# Structural basis of the P4B ATPase lipid flippase activity

**DOI:** 10.1101/2021.04.28.441804

**Authors:** Lin Bai, Bhawik K. Jain, Qinglong You, H. Diessel Duan, Todd R. Graham, Huilin Li

## Abstract

P4 ATPases are lipid flippases that are phylogenetically grouped into P4A, P4B and P4C clades. The P4A ATPases are heterodimers composed of a catalytic α-subunit and accessory β-subunit, and the structures of several heterodimeric flippases have been reported. The *S. cerevisiae* Neo1 and its orthologs represent the P4B ATPases, which function as monomeric flippases without a β-subunit. It has been unclear whether monomeric flippases retain the architecture and transport mechanism of the dimeric flippases. Here we report the first structure of a P4B ATPase, Neo1, in its E1-ATP, E2P-transition, and E2P states. The structure reveals a conserved architecture as well as highly similar functional intermediate states relative to dimeric flippases. Consistently, structure-guided mutagenesis of residues in the proposed substrate translocation path disrupted Neo1’s ability to establish membrane asymmetry. These observations indicate that evolutionarily distant P4 ATPases use a structurally conserved mechanism for substrate transport.

## INTRODUCTION

The phospholipid bilayer of biological membranes provides a fundamental barrier that encloses the cell and most internal organelles, while integral membrane proteins allow for the selective passage of ions and polar molecules. The lipid organization of many membranes is asymmetric. For example, the plasma membrane contains more phosphatidylethanolamine (PE) and phosphatidylserine (PS) in its cytosolic leaflet, and more phosphatidylcholine (PC) and sphingolipids in the extracellular leaflet ^1–3^. Phospholipid asymmetry is crucial for many cellular processes, including vesicular transport, signal transduction, cell motility, cell polarity, cell division, and the transport of ions and other molecules across the bilayer ^1,4–7^. Phospholipid asymmetry is regulated by three types of transporters: scramblases, floppases, and flippases ^8,9^. Scramblases catalyze nonselective, bidirectional, trans-bilayer movement of the phospholipid down their concentration gradients. Floppases belong to the ABC transporter family and translocate phospholipids to the extracellular leaflet of membranes. Flippases belong to the type IV P-type ATPase (P4 ATPase) family and selectively transport phospholipids to the cytosolic leaflet of membranes.

P-type ATPases form ATP-dependent phosphorylated intermediates during their transport of substrates across biological membranes ^10^. Based on sequence homology, P-type ATPases are divided into five subclasses (P1 to P5) ^11–14^, and each subclass typically transports different substrates. The P1, P2, and P3 ATPases mainly transport cations (e.g., the Ca^2+^-ATPase, Na^+^/K^+^-ATPase, and the proton pump) or heavy metals (e.g. the copper transporter). In stark contrast, the substrates of P4 ATPases are bulky phospholipid molecules, and these transporters are further divided into P4A, P4B, and P4C clades ^15^. The P5 ATPases transport polyamines or transmembrane helices ^16–19^. Most P-type ATPases have a conserved architecture consisting of a transmembrane domain (TMD), an actuator domain (A-domain), a nucleotide-binding domain (N-domain), and a phosphorylation domain (P-domain) ^12^. According to the structural studies of the most well-understood P2 ATPases, P-type ATPases transport their substrate through a cyclic transition of E1–E1P–E2P–E2 states, which was called the “Post–Albers” model ^20–24^.

*Saccharomyces cerevisiae* has five P4 ATPases—Drs2, Neo1, Dnf1, Dnf2 and Dnf3, each having a different subcellular location and substrate specificity—whereas humans have 14 ^25–27^. Notably, most P4 ATPases transport phospholipids as a heterodimeric complex, with the major catalytic subunit referred to as the α-subunit and the partner protein as the β-subunit. The β-subunit is essential for the stability and trafficking of the flippase, as well as for substrate binding ^28–31^. For P4A ATPases, the α-subunit makes extensive contacts with the β-subunit on the extracellular side of the membrane, within the membrane, and on the cytosolic side. These αβ interactions are thought to be crucial for the distinct conformation dynamics of P4 ATPases that control lipid translocation. However, the yeast Neo1 and its related human proteins (ATP9A and ATP9B) are monomeric, and the structural basis for how they function without the accessory β-subunit has been unclear.

Neo1 was first identified as a neomycin-resistance-1 gene (*NEO1*) that prevented aminoglycoside toxicity when overexpressed ^32^. Neo1 is essential for cell growth and is found in the Golgi and endosomes, where it regulates membrane trafficking ^33–36^. Although the flippase activity of Neo1 has not been demonstrated with purified protein, temperature-sensitive alleles of *NEO1* cause a loss of PE and PS asymmetry in the plasma membrane ^37^. However, Neo1 plays a more substantial role in preventing exposure of PE than PS, suggesting that Neo1 mainly transports PE to the cytosolic side ^37^. Although it lacks a β-subunit, Neo1 is regulated by Dop1 (*Dopey* ortholog), Mon2 (which is a relative of large Arf guanine nucleotide exchange factors), the small GTPase Arl1, and Any1 ^33,38–41^. In fact, Neo1, Mon2, Arl1, and Dop1 assemble into a membrane remodeling complex ^38,39^. All these proteins interact with Neo1 and are crucial for yeast growth but have few known functions. The disruption of *ANY1*, which encodes a PQ-loop membrane protein, can rescue the growth deficiency of a *Δneo1* or *Δdrs2* strain ^37,41,42^.

Recently, several P4A ATPase structures have been reported, including the *S. cerevisiae* Drs2–Cdc50 ^43,44^, Dnf1–Lem3 ^31^, Dnf2–Lem3 ^31^, the *C. thermophilum* Dnf1–Cdc50 ^45^, the human ATP8A1–CDC50A ^46^, and ATP11C-CDC50A ^47^. These structural studies revealed a conserved architecture among the P4 ATPases, including the 10-helix TMD, the cytosolic A domain, N domain, and P domains, and a conserved ATP-dependent lipid transport cycle. Several of the P4A ATPase structures contain substrate lipid bound in an exofacial “entry site” formed from residues that had previously been implicated in substrate recognition through mutagenesis studies ^31,47–49^. A second substrate lipid-binding site on the cytosolic side of the membrane, an exit site, has been identified only in the Dnf1 and Dnf2 structures ^31,45^. Surprisingly, this large exit gate extends 10 Å out of the bilayer, causing membrane to dimple into the cytosol, and it uses an Arg residue from the N-terminal cytosolic domain of the β-subunit to help coordinate the lipid headgroup. However, it has been unclear whether the monomeric lipid flippase Neo1 functions by a similar mechanism or whether the cytosolically exposed exit gate is a conserved aspect of the substrate translocation path for P4 ATPases that lack a β-subunit.

An important question is how the substrate specificity is achieved for the P4B ATPases. Inactivation of Neo1 perturbs the PS and PE asymmetry of the plasma membrane, but whether or not Neo1 interacts directly with these lipids as transport substrates is unclear. By comparison, Drs2–Cdc50 primarily transport PS, while Dnf1–Lem3/Dnf2–Lem3 transports glucosylceramide (GlcCer), PC, and PE ^50,51^. The lipid substrate specificity of P4A ATPases appears to be defined by the physiochemical properties of these substrate-binding pockets ^52^, but how these P4 ATPases select their substrate is only partially understood. To address these questions, we purified the yeast Neo1; examined its substrate-stimulated, in vitro ATP hydrolysis activity; determined cryo-EM structures of Neo1 in three intermediate states; and performed an extensive structure-guided mutagenesis and functional assays. Our work identifies unique residues responsible for substrate specificity and provides insights into the conservation of the cytosolic binding site despite the lack of a β-subunit.

## RESULTS AND DISCUSSION

### 1. Neo1 ATPase activity is stimulated by PE and PS

We overexpressed Neo1 in a yeast strain using a multicopy plasmid with a strong GAP promoter and an N-terminal triple FLAG tag. The detergent dodecyl maltoside (DDM) was used to solubilize the membrane and we purified Neo1 with an anti-FLAG affinity column followed by size-exclusion chromatography, during which DDM was replaced by lauryl maltose neopentyl glycol (LMNG) and cholesteryl hydrogen succinate (CHS) to stabilize the membrane protein (**Fig. 1a-b**). Neo1 has been implicated in the transport of PS and PE in vivo ^37^. We used substrate-stimulated ATP hydrolysis activity as an indicator to investigate whether the purified protein was active and whether PE and PS directly interacted with Neo1. Dnf1–Lem3 displays a preference for lyso-phospholipid substrate ^49^, so we also tested lyso-PS as a potential substrate. We found that the Neo1 ATPase activity was stimulated by PE, PS, or lyso-PS, each at 0.1 mM (**Fig. 1c**). Furthermore, we demonstrated that the ATPase activity of Neo1 increased as the substrate concentration increased (**Fig. 1d**). Prior studies of the influence of Neo1 on membrane asymmetry suggested PE was the preferred Neo1 substrate ^34^ but both PS and PE effectively stimulated ATPase activity. These data showed that the purified Neo1 was active and that the PS, PE, and lyso-PS are very likely native substrates of the flippase.

**Fig. 1.**
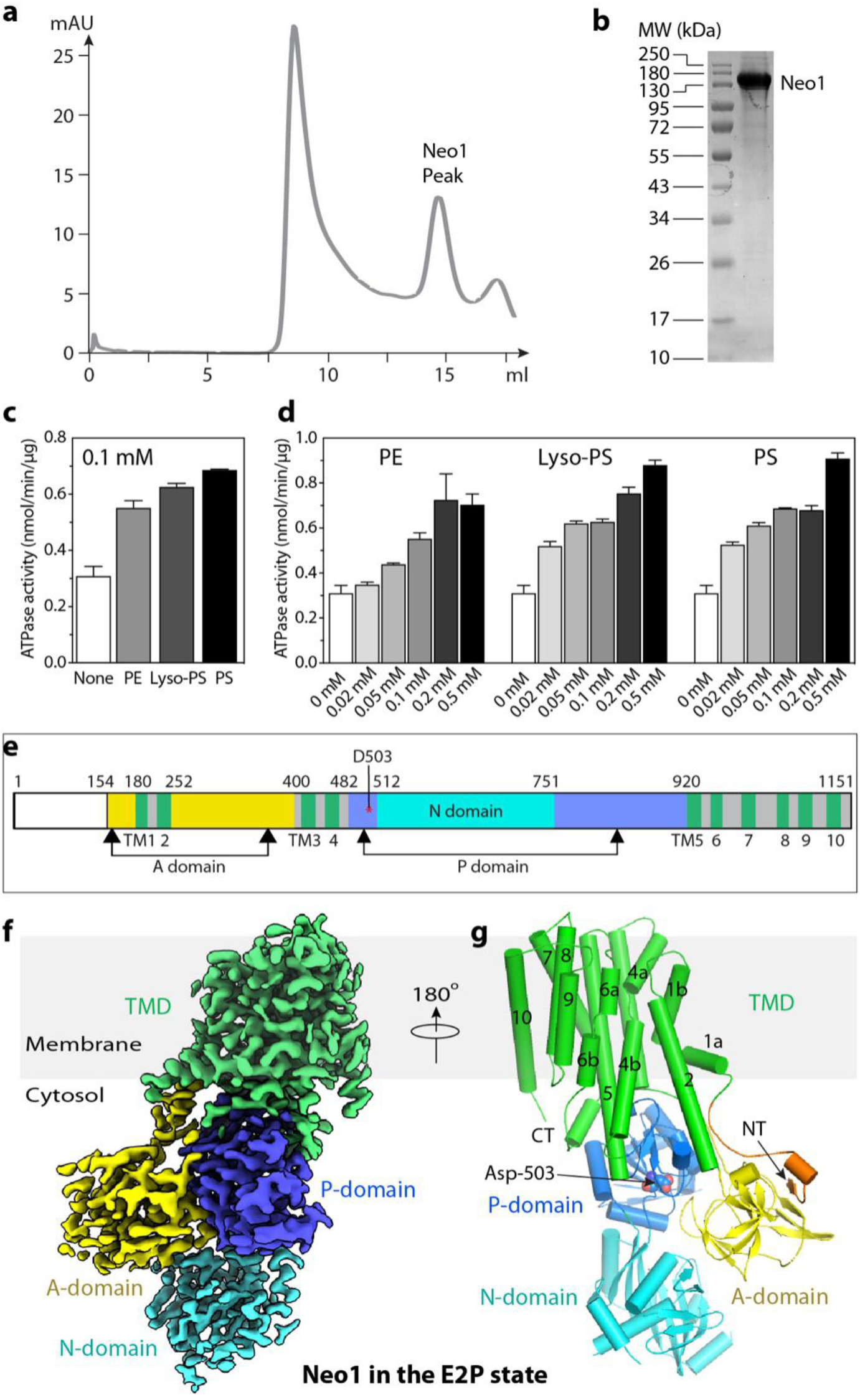
Cryo-EM of the purified and active Neo1. **(a)** Gel filtration profile of Neo1 in detergent. (**b**) SDS-PAGE gel of the purified Neo1. (**c**) Substrate stimulated the ATP hydrolysis activity of Neo1. Substrates were PE, PS, and lyso-PS, all at 0.1 mM concentration. (**d**) Lipid concentration-dependent ATPase activity of Neo1. Data points in (c-d) represent the mean ± SD in triplicate. (**e**) The Neo1 domain map. (**f**) Cryo-EM 3D map of Neo1 in the E2P in the front view. The major domains are labeled in different colors. **g**) Atomic model of Neo1 in the E2P state in cartoon shown in the back view and colored as in panel **f**. The phosphate acceptor residue Asp-503 in the P domain is shown in spheres. The ordered N-terminal β-strand and short α-helix are in orange.

### 2. Cryo-EM structure of Neo1 in the E2P state

We performed single-particle cryo-EM on purified samples either directly or incubated with different inhibitors to capture structures in different states. We obtained cryo-EM 3D maps of Neo1 in the E1-ATP state stabilized by AMPPCP at 5.6-Å resolution, in the E2P-transition state stabilized by AlF_4^−^_ at 3.1-Å resolution, and in the E2P state as stabilized by BeF_3^−^_ at 3.2-Å resolution (**Supplemental Table 1, Supplemental Figures 1-4**). We found that 3D maps of Neo1 in the E2P-transition and E2P states were superimposable (**Supplemental Figure 5**). This is consistent with the previous observation that the Dnf2–Lem3 structures in the two states are nearly indistinguishable ^31^. We first built atomic models into the 3.2-Å resolution map in the E2P state and into the 3.1-Å resolution map in the E2P-transition state. The 3D maps had excellent main-chain connectivity and side-chain densities except for four disordered loops: the N-terminal tail (residues 1-153), the N-domain loop (542-559), the P-domain loop (804-821), and the C-terminal tail (1142-1151) (**Fig. 1D**). We next used the E2P structure to help build an atomic model for the lower-resolution 3D map of Neo1 in the E1-ATP state. All these models fit well with the 3D maps and were refined to good statistics (**Supplemental Table 1**).

As shown in the E2P state structure, Neo1 contains all conserved P-type ATPase domains: a 10-TM helix (TMH1-10) TMD; the cytosolic A domain inserted between TMH2 and TMH3; and the cytosolic P and N domains that are both inserted between TMH4 and TMH5 (**Fig. 1d-e**). The Neo1 structure aligns well in both TMD and soluble domains with the α-subunits of heterodimeric P4 ATPases Dnf1–Lem3, ATP8A1–CDC50, ATP11C–CDC50, and Drs2–Cdc50, (**Supplemental Figure 6**). The most distinct feature of Neo1 is the very short extracellular loops (ECLs) (**Fig. 1f-g**), as they do not need to bind and coordinate a β-subunit. Note that TMH1, TMH4, and TMH6 of Neo1 are kinked just as they are in other P-type ATPases whose structures are known. This conserved structural feature is essential for the substrate transporting activity. The Neo1 TMH4 kink is enabled by the highly conserved Pro-456.

The N- and C-terminal peptides play regulatory roles and share low sequence homology among the P4 ATPases. For examples, the N-terminal peptides of Dnf1 and Dnf2 contain phosphorylation sites that are essential for activation ^53^; the C-terminal peptides of the yeast Drs2 and the corresponding human homolog ATP8A1 are autoinhibitory ^43,44,46,54,55^. The N-terminal peptide of Neo1 interacts with the cargo-selective sorting nexin Snx3 to mediate Neo1 trafficking ^35^. The N-terminal peptide preceding the TMH1 is 183 residues long, of which the first 153, containing the Snx3 binding site, were disordered in our structure. The 30-residue ordered region of the N-terminal peptide (Glu-154 through Val-183) formed a short β-strand and a short α-helix that binds to the regulatory A domain (**Fig. 1g**). Unlike other P4 ATPases, the Neo1 C-terminal peptide following the TMH10 (Tyr-1142 to Pro-1151) is short and disordered. Therefore, it is unlikely that the Neo1 C-terminal tail is an autoinhibitory domain comparable to Drs2 or ATP8A1.

### 3. Neo1 structure in the E1-ATP state and the conserved ATP-dependent transport mechanism

In the 5.6-Å resolution 3D map of Neo1 in the E1-ATP state determined in the presence of 1 mM AMPPCP (a nonhydrolyzable ATP analog), the AMPPCP density is clear and the molecule stabilizes the interface between the N and P domains (**Fig. 2a**). Comparison of the Neo1 structures in the E1-ATP and the E2P states shows that TMHs 3-10 and the P domain are superimposable, whereas TMHs 1-2 have moved slightly, while the N and A domains have undergone dramatic conformational changes (**Fig. 2b**). In the E1-ATP structure, the N domain is packed tightly on P domain through the AMPPCP ligand. The A domain is also ordered, although it has no interaction with the P domain and a very weak interaction with a short helix-turn-helix motif (Lys704 to Leu736) in the N domain. The long TMH2 bends upwards at Lys-236 away from the P domain. Transitioning from the E1-ATP state to the E2P state, ATP is hydrolyzed, ADP is released from the N domain, and the phosphate is transferred onto Asp-503 of the P domain. The phosphorylated aspartate is mimicked by the BeF_3^−^_ moiety in our Neo1 structure. To enter into the E2P state, both the N and A domains swing away from the membrane and pack tightly together. Accompanying this transition, the TMH2 becomes straightened, pointing downwards to allow the A domain movement. These large conformational changes are likely propagated to the substrate binding sites and alter the substrate binding affinity, leading to the transport of a lipid molecule across the membrane. Conversely, the transition from E2P →E2→E1-ATP should close up the exit site and release substrate into the cytosolic leaflet.

**Fig. 2.**
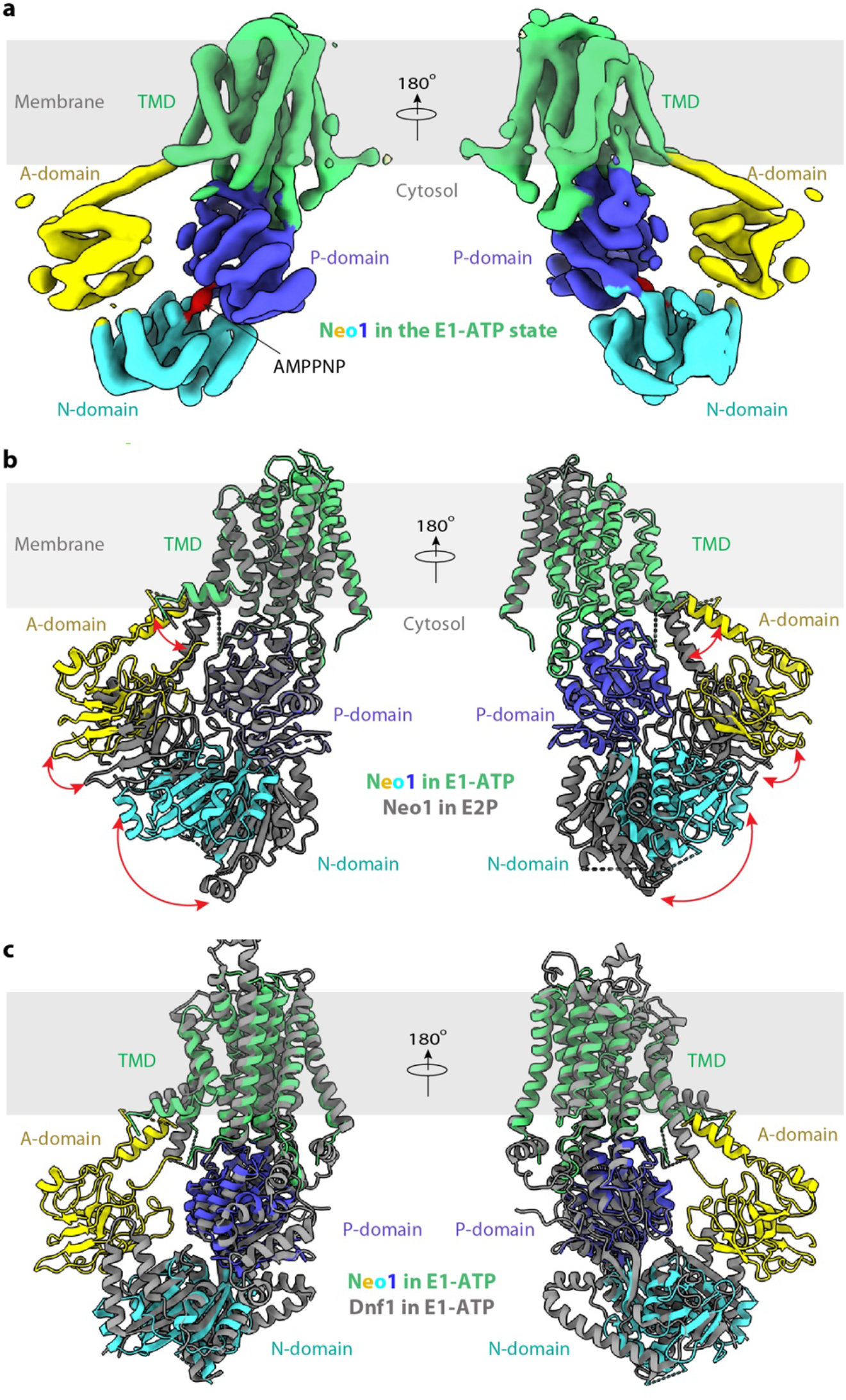
Structure of the *S. cerevisiae* Neo1 in the E1-ATP state. **a**) Cryo-EM 3D map of Neo1 in the E1-ATP, in front (left) and back (right) views. The major domains are labeled in different colors. The AMPPNP density is red. **b**) Structural comparison between Neo1 in the E1-ATP state (colors) and in the E2P state (grey). **c**) Comparison of Neo1 (colors) and Dnf1 (grey) in the E1-ATP state.

We next compared the Neo1 E1-ATP structure with the published structure of Dnf1–Lem3 in the same state and found that the TMD and the cytosolic domains are positioned similarly (**Fig. 2c**). Furthermore, the conformation and the substrate transport path of Neo1 in the E2P state highly resemble those of the Drs2, Dnf1, ATP8A1, and ATP11C flippases in the same E2P state (**Supplemental Figure 6**). The lack of TMH3-10 movement through the P4 ATPase transport cycle was proposed to be conferred by the tightly associated β-subunit ^46^, but Neo1 retains this property in the absence of a stabilizing β-subunit. This observation strongly suggests that all P4-ATPases—whether functioning as a single subunit in the case of Neo1 or with a partner β-subunit in the majority of cases—employ a conserved substrate transport mechanism. A recent functional study of Neo1 appears to support such a conserved transport mechanism for the substrate entry site ^56^.

### 4. The substrate transport path and the substrate specificity of Neo1

Structures of six P4 ATPases have been reported so far; four are in complex with substrate lipids ^31,43–47^ (**Supplemental Figure 6**). These structures reveal a conserved flippase architecture, similar conformational changes as the enzymes go through the transport cycle, and a very similar substrate translocation path, despite having different substrate specificities. A structure-based homolog search revealed that Neo1 in the E2P state aligned well with all known P4 ATPase structures in that state; but the best match was with Dnf1 (**Fig. 3a**). Yet the superposition of Neo1 and Dnf1 revealed a major difference between these flippases: the ECL2, 4, and 5 of Dnf1, which are primarily responsible for binding the β-subunit Lem3, are much longer than those in Neo1 (**Fig. 3a-b**). These extracellular loops are similar in length between Neo1 and cation transporters. This observation led us to hypothesize that P4A ATPases evolved into two-subunit ATPases by the lengthening of ECL2,4 and 5, thereby acquiring the ability to bind a b-subunit.

**Fig. 3.**
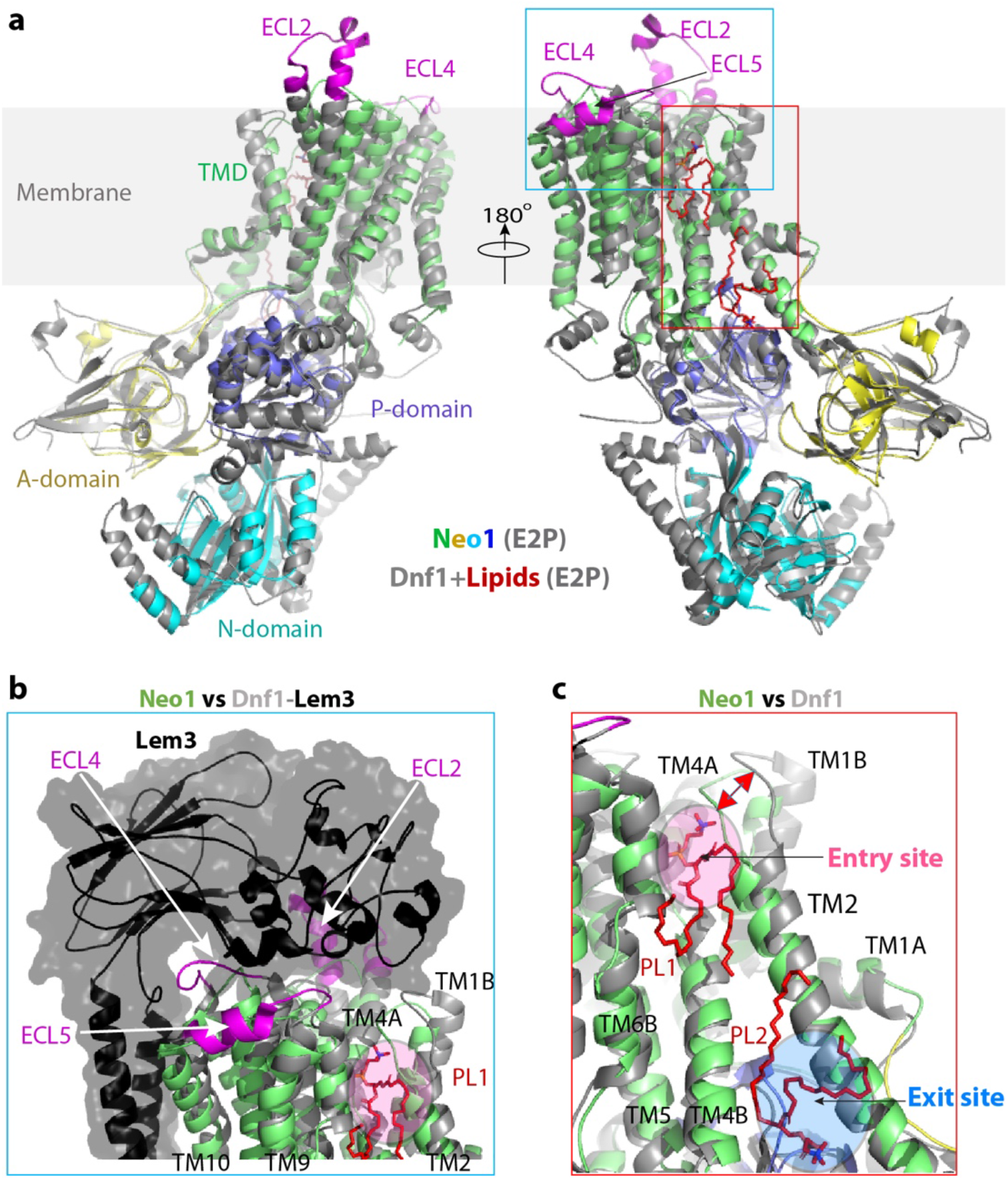
Structural comparison between Neo1 and Dnf1 in the E2P state. **a**) Lipids in the Dnf1 structure are shown as red sticks. The PL binding sites are demarcated by the red rectangle and enlarged in (c). The cyan square marks the β-subunit interacting top region of the Dnf1. All flippases have five extracellular loops (ECLs) connecting TM1 and 2, TM3 and 4, and TM5 and 6, TM7 and 8, and TM9 and 10. ECL2, 4, and 5 of Dnf1 (magenta) are much longer than those of Neo1. **b**) The extended ECL2, 4, and 5 of Dnf1 insert into and tightly bind the β-subunit Lem3; the corresponding loops of Neo1 are short and Neo1 lacks a β-subunit. **c**) Enlarged view of the substrate transport path of Neo1 and Dnf1. The pink and blue ovals mark the lipid entry and exit sites. The double-headed red arrow marks the 7-Å outward movement of TM1B and associated ECL1 relative to Neo1, due to the presence of a lipid at the entry site in Dnf1.

Previous functional studies suggested that Neo1 transported both PE and PS, but how Neo1 recognizes these lipids was unknown ^37^. We did not capture any endogenous lipid molecules in the Neo1 substrate sites. However, the Dnf1 structure contained phospholipid molecules in both the substrate entry and exit sites. By aligning the Neo1 and Dnf1 structures, we narrowed down the substrate translocation path of Neo1 to the back side of the ATPase between TMH1-4 and TMH6 (**Fig. 3a, c**). This putative substrate path can be divided into three sections: the top (lumenal) substrate entry site composed of several polar residues (Gln-209, Ser-221, Tyr-222, Ser-452, and Thr-453); the middle transport path between lipid-binding sites that contains the hydrophobic gate (Phe-202, Leu-226, Val-229, Val-457, and Val-461) and flanking polar residues (Gln-193, Thr-233, Arg-460, and Asp-464); and the bottom (cytosolic) substrate exit site lined by many polar residues (Lys-236, Asp-240, Gln-243, Arg-247, Ser-468, Glu-475, and Ser-488) (**Fig. 4a**). To identify key residues responsible for Neo1 substrate-transporting activity and specificity, we next carried out an extensive structure-guided mutagenesis and functional assays (**Fig. 4b, Supplemental Figures 7-8**). Most of these residues were mutated to either alanine or to residues present at the same position in other P4 ATPases.

**Fig. 4.**
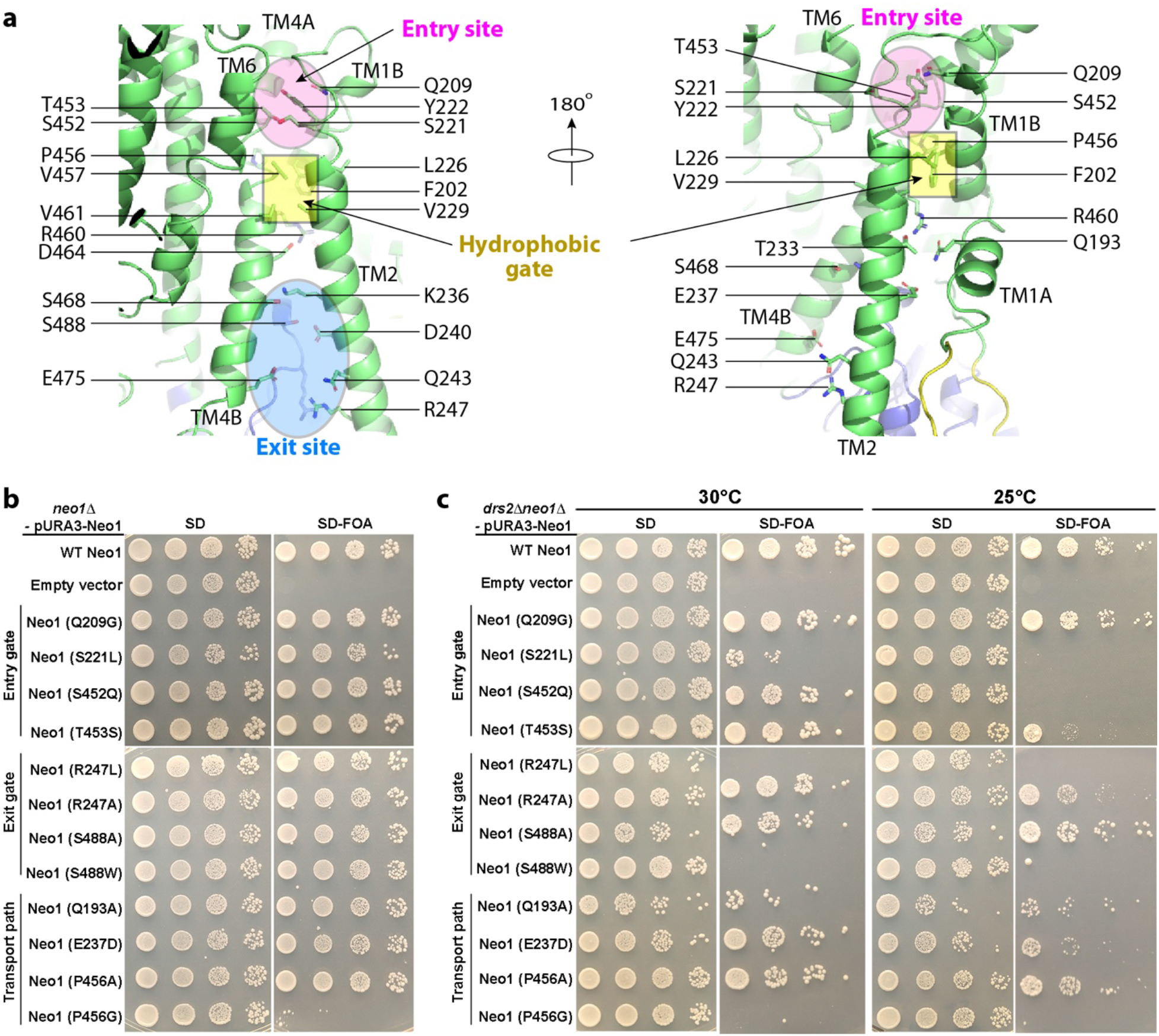
Mutagenesis of the substrate path. **a**) Putative substrate transport path of Neo1. Cartoon view of the substrate-transporting channel in the Neo1 structure with the residues lining the path shown as sticks. Growth phenotypes of *neo1*Δ (**b**) and *neo1*Δ *drs2*Δ (**c**) strains expressing Neo1 mutant variants. Strains on the SD medium expressed both WT and mutant variants of Neo1, but SD+5FOA medium eliminated WT Neo1 expressed from the *URA3*-marked plasmid. Colony size on the 5-FOA medium indicates how well the Neo1 variants support growth.

Neo1 is essential for cell viability, and so we used a plasmid shuffle assay to test whether Neo1 variants could support viability of a *neo1*Δ strain (**Fig. 4b**, **Supplemental Figure 7**). The parental *neo1*Δ strain harboring wild-type (WT) *NEO1* on a *URA3*-marked plasmid was transformed with *HIS3*-marked plasmids harboring either WT Neo1, an empty vector, or the Neo1 mutants. The strains were replica-plated on SD medium to select for both plasmids, or on SD plus 5-FOA to counter-select against the *URA3* plasmids. Because *neo1*Δ cells are inviable, the empty vector control strain failed to grow on 5-FOA, while the WT copy of Neo1 fully supported growth. We observed that all of the Neo1 mutants supported viability comparably to WT Neo1 except for Neo1 P456G, which failed to support growth (**Fig. 4b**). Pro-456 is near the center of M4, and nearly all P-type ATPases have a proline in this position that is crucial for unwinding of the M4 helix. We found that Neo1 P456G totally abolished the Neo1 activity in vivo. Surprisingly, Neo1 P456A fully supported viability, suggesting that an alanine at this position is compatible with a distorted helix and that maintenance of hydrophobicity in this region is essential (**Fig, 4b**).

Previous studies indicated that there is partial functional redundancy between Neo1 and Drs2. Therefore, we also tested the Neo1 mutants for their ability to support growth of a *drs2*Δ *neo1*Δ double mutant over a range of temperatures, which provides a more sensitive assay for loss of Neo1 function (**Fig. 4c, Supplemental Figure 8**). *drs2*Δ single mutants grow well at 30 °C, but are cold-sensitive and grow slowly at 25 °C and fail to grow at 20 °C. We found that the exit-site mutations Neo1 S488W and Neo1 R247L were lethal when combined with *drs2*Δ and failed to grow at 30 °*C*. Many of the other Neo1 entry-site and transport-pathway mutations substantially reduced growth in this background relative to WT Neo1. These results indicate that the Neo1 variants represent an allelic series, ranging from complete loss of function (P456G) to apparent WT activity, with respect to cell viability (P456G < S488W = R247L < S221L < Q193A < S452Q < E237D < R247A < P456A = T453S < S488A = Q209G).

Neo1 is primarily found in the Golgi, and a loss of its flippase activity should increase substrate lipids in the luminal leaflet of the Golgi and their ultimate exposure in the extracellular leaflet of the plasma membrane. Therefore, to test the influence of Neo1 mutations on substrate transport, we employed toxin sensitivity assays to measure the exposure of PS and PE on the plasma membrane extracellular surface (**Fig. 5a**). Papuamide A (Pap A) is potent cytotoxic agent that binds specifically to PS, and duramycin binds specifically to PE. Both toxins produce pores in the membrane that kill sensitive cells on which their respective targets are exposed. Wild-type yeast cells are relatively resistant to Pap A and duramycin because most of their target lipids are restricted to the inner cytosolic leaflet of the plasma membrane.

**Figure 5:**
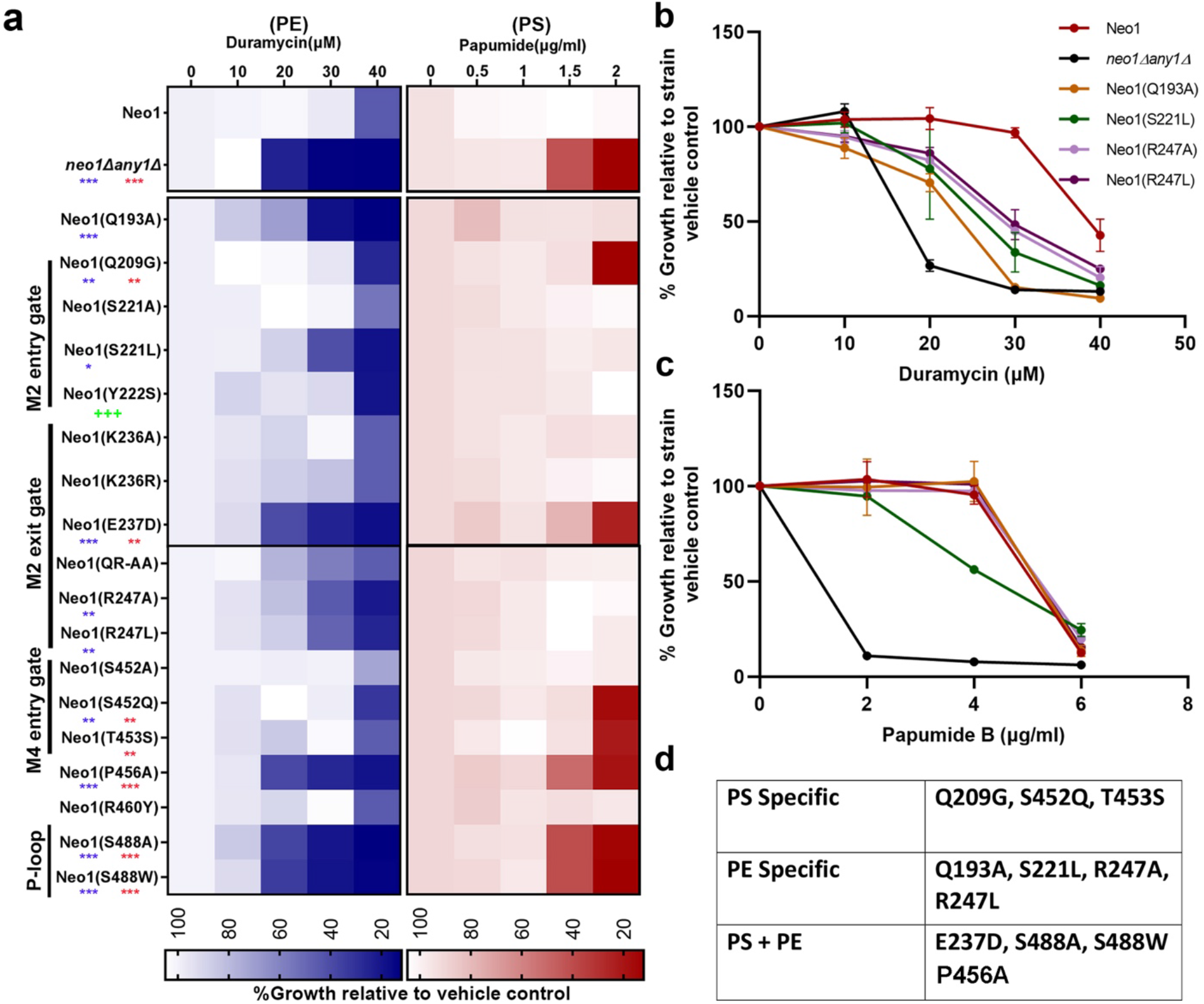
Neo1 substrate pathway mutations disrupt plasma membrane asymmetry. **a)** Duramycin and papuamide sensitivity of *neo1*Δ strains expressing the indicated Neo1 mutant variants relative to *neo1*Δ expressing WT Neo1 and *neo1 any1*Δ (a viable strain displaying a complete loss of function for Neo1’s influence on membrane asymmetry). The blue heat map displays duramycin dose responses; growth inhibition at 20-30 μM indicates aberrant exposure of PE. The red heat map displays papuamide A dose responses; growth inhibition at 1.5–2 μg/mL indicates aberrant exposure of PS. **\*** p < 0.05, **\*\*** p < 0.01, **\*\*\*** p < 0.001 (red for papuamide and blue for duramycin). ^+++^Neo1 (Y222S) is a gain-of-function mutation that weakly suppresses *drs2*Δ. **b-c**) Neo1 mutants hypersensitive to only duramycin in (**a**) were tested at higher concentrations of papuamide (up to 6 μg/mL) (n = 3, +/−SD). **d)** Table summarizing the effects of Neo1 mutations on membrane asymmetry.

The *neo1*Δ strain expressing a WT copy of Neo1 was resistant to duramycin and papuamide relative to the *neo1*Δ*any1*Δ strain, which exposed both lipids and was hypersensitive to both toxins. The *any1*Δ mutation suppresses *neo1*Δ lethality but does not suppress the loss of PS and PE asymmetry caused by loss of Neo1 function ^42^. Relative to these two control strains, many of the Neo1 mutants displayed a significant loss of membrane asymmetry. We further explored the Neo1 mutants that had PE exposure and no apparent PS exposure by using higher concentrations of papuamide to assess how well they maintained PS asymmetry (**Fig. 5b-c**). Neo1-S221L displayed a slight hypersensitivity to papuamide, but the rest of the mutants showed sensitivity indistinguishable from that of the WT.

The Neo1 mutants fell into three classes with regard to substrate recognition (**Fig. 5d**). PS-specific mutants that expose PS but maintain PE asymmetry (Q209G, S452Q, and T453S), PE-specific mutants that only expose PE (Q193A, S221L, R247A, and R247L), and mutants that expose both lipids. These substrate-specific, separation-of-function mutations strongly imply that the targeted residues are directly involved in substrate binding. For example, entry-site residues known to bind PS in ATP8A1–CDC50 correspond to the PS-specific Neo1 mutants. This is the first study to define residues crucial for PE recognition in any P4 ATPase, and these residues are in the entry site (S221), transport path (Q193), and exit site (R247). Neo1 mutations that disrupt PE transport generally caused the strongest growth defect when combined with *drs2*Δ, although the correlation between growth defects and loss of membrane asymmetry is imperfect (**Figs. 4-5**).

Mutations that decrease transport of both PS and PE are not likely causing misfolding of Neo1 because these alleles fully support the growth of *neo1*Δ cells (**Fig. 4b**). We tagged such Neo1 variants with GFP and examined their localization relative to mKate-Aur1, a Golgi marker. Mutations that significantly alter structure of membrane proteins in the secretory pathway typically cause retention of the mutant protein in the endoplasmic reticulum (ER) ^31,57^. However, the Neo1 variants we tested localized normally to the Golgi complex (**Supplemental Figure 9)**. Thus, these mutations likely cause a loss of either substrate recognition or transport rather than a structural defect.

### 5. A comparison of the substrate-binding sites of lipid flippases

According to previous functional studies, each type of P4 ATPase may transport several different lipid substrates, but the preference is usually not equal ^34,50,51^. We hypothesize that the substrate preference is likely determined by the chemical and electrophysiological properties of the substrate entry site. Having defined the substrate-binding sites in Neo1 via mutagenesis and functional assays, we compared the substrate entry sites of Neo1, Dnf1, Drs2, ATP8A1, and ATP11C (**Fig. 6a**), which are all in the back surface and surrounded by TMH1, TMH2, TMH4, and TMH6. We found that they all contained polar and non-charged residues such as Ser, Thr, Asn, and Gln. Structure-based sequence alignment showed that key residues at structurally equivalent positions were not well conserved in the primary sequence (**Fig. 6b**). The Neo1 functionally essential dipeptide motif 209-QA-210 in TMH1 is analogous to GA in Dnf1 and Dnf2, QQ in Drs2 and ATP8A1, and the single residue Q in ATP11C. The Neo1 dipeptide motif 221-SY-222 in TMH2 is analogous to TT in Drs2 and ATP8A1, TS in ATP11C, LS in Dnf1, and FA in Dnf2. The Neo1 dipeptide motif 452-ST-453 becomes NN in ATP8A1, SN in Drs2, NF in ATP11C, and QS in both Dnf1 and Dnf2. Finally, the Neo1 residue Ala (A978) becomes an asparagine in all the other P4 ATPases compared here. The idea that these residues in the entry sites of the P4 ATPases dictate the lipid-transporting preference is underscored by previous mutational studies. In one study, mutations of the TMH1 key motif GA and the TMH4 motif QS of Dnf1/Dnf2 were shown to alter their activity for PC, PE, and GlcCer, and mutation of the TMH6 key reside N1226 of Dnf1 abolished the transport of all three substrates ^51^. In another study, the TMH1 key motif QQ of Drs2 was shown to influence the specificity for PS transport ^48^.

**Fig. 6.**
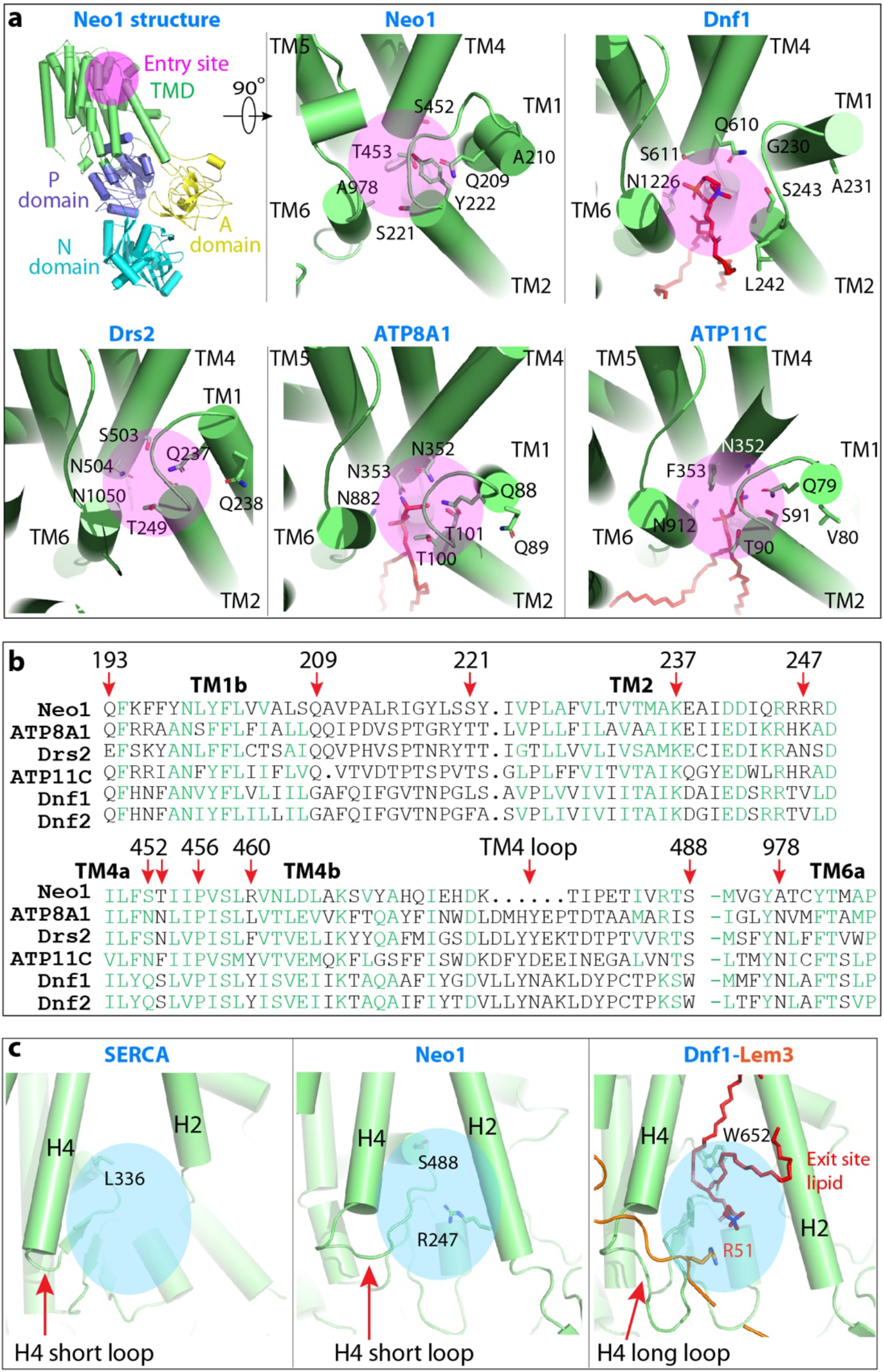
Structural comparison of substrate entry and exit sites in Neo1 and other P4 ATPases. **a**) Views of substrate entry sites in Neo1, Dnf1 (PDB ID 7KYC), Drs2 (PDB ID 6PSY), ATP8A1 (PDB ID 6K7M), and ATP11C (PDB ID 7BSV). Key residues and lipids (if present) are shown as sticks. The entry sites are highlighted by a magenta circle. **b**) Structure-based sequence alignment of Neo1, Dnf1, Dnf2, Drs2, ATP8A1, and ATP11C. Key residues involved in substrate lipid binding are highlighted by red arrows. Conserved residues are shown in green. **c**) Comparison of substrate exit sites in SERCA (PDB ID 2AGV), Neo1, and Dnf1–Lem3 (PDB ID 7KYC). The blue circles mark the substrate exit sites.

The substrate-binding exit site initially defined in Dnf1–Lem3 and Dnf2–Lem3 was unanticipated, because it extends substantially out of the membrane plane into the cytosolic domains of these P4A ATPases. This exit site is structurally conserved in all of the P4 ATPase structures (**Supplemental Figure 6**). It is formed from cytosolic extensions of TMH2 and TMH4 and a loop emanating from TMH4 that forms the membrane proximal region of the P domain. We show in the current study that this cytosolically exposed lipid-binding site is functionally required for substrate transport in Neo1, a P4B ATPase. Comparison of the exit sites suggests a mechanism for how Neo1 functions without a β-subunit. The most membrane-distal residue forming the exit site in Dnf1–Lem3 is R51 in the cytosolic N-terminal tail of Lem3, analogous to R247 in Neo1 on the TMH2 side of the cleft (**Fig. 6c**). The β-subunit N-terminal tail interacts with the loop from TMH4 that bends back towards the membrane to form part of the exit site (W652 in Dnf1), before starting the P-domain where the phosphorylated Asp resides. This TMH4 loop is much longer in the P4A ATPases than in the P4B ATPases and is nearly absent in P2 ATPases (e.g., SERCA) (**Supplemental Figure 10**).

In summary, we have determined the first structure of a single-subunit lipid flippase from any species. The structure suggests that P4B ATPases such as Neo1 arose first from cation transporters and gave rise to the P4A group with their new β-subunit requirement. The overall architecture and the substrate entry site of Neo1 are remarkably conserved in structure but, interestingly, not in amino acid sequence. Key differences between the P4A ATPases and P4B ATPases reflect regions important for β-subunit interaction, including the shorter extracellular loops in Neo1, a shorter TMH4 cytosolic loop, and the ability to form the exit site in the absence of the β-subunit. Our extensive mutational studies identified key residues responsible for Neo1’s substrate preference. Structural comparison of Neo1 with other P4 ATPases revealed that the basic ATP-dependent lipid transport mechanism is highly conserved throughout evolution. The capability and preference of the ATPases for transporting a multitude of lipid species are likely conferred by varying the amino acids of the key motifs in the otherwise structurally similar substrate entry and exit sites. Our work has advanced the mechanistic understanding of a large and often essential family of the P4 ATPases.

## MATERIALS AND METHODS

### Expression and purification of Neo1

*S. cerevesiae* cells were grown in 2 L SD-H medium for about 20 h, then transferred to 18 L of YPD medium for another 12 h before harvest. Cells were resuspended in lysis buffer (20 mM Tris-HCl, pH 7.4, 0.2 M sorbitol, 50 mM potassium acetate, 2 mM EDTA, and 1 mM phenylmethylsulfonyl fluoride [PMSF]) and then lysed using a French press at 15,000 psi. The lysate was centrifuged at 10,000 × *g* for 30 min at 4 °C. The supernatant was collected and centrifuged at 100,000 × *g* for 60 min at 4 °C. The membrane pellet was collected and then resuspended in buffer A containing 10% glycerol, 20 mM Tris-HCl (pH 7.4), 1% DDM, 0.1% CHS, 0.5 M NaCl, 1 mM MgCl_2_, 1 mM MnCl_2_, 1 mM EDTA, and 1 mM PMSF. After incubation for 30 min at 4 °C, the mixture was centrifuged for 30 min at 120,000 × *g* to remove insoluble membrane. The supernatant was mixed with pre-equilibrated anti-FLAG (M2) affinity gel at 4 °C overnight with shaking. The affinity gel was then collected and washed three times in buffer B (20 mM HEPES, pH 7.4, 150 mM NaCl, 0.01% LMNG, 0.001% CHS, and 1 mM MgCl_2_). The proteins were eluted with buffer B containing 0.15 mg/mL 3×FLAG peptide and was further purified in a Superose 6 10/300 Increase gel filtration column in buffer C (20 mM HEPES, pH 7.4, 150 mM NaCl, 0.0025% lauryl maltose neopentyl glycol (LMNG), 0.00025% cholesteryl hydrogen succinate (CHS), and 1 mM MgCl_2_). Finally, the purified proteins were assessed by SDS-PAGE gel and concentrated for cryo-EM analysis. Approximately 20 μg of Neo1 can be purified from 18 L of cells.

### ATP hydrolysis assay

The ATPase activity assays were performed using BIOMOL Green Reagent (Enzo Life Sciences, Inc.) to measure released inorganic phosphate. The lipids phosphatidylethanolamine, lyso-phosphatidylserine, and phosphatidylserine were solubilized with 20 mM sodium cholate in 20 mM HEPES, pH 7.4, 150 mM NaCl. Each reaction contained 0.025 mg/mL protein, 0.003% LMNG, 0.0003% CHS, 20 mM HEPES, pH 7.4, 150 mM NaCl, 10 mM MgCl_2_, and 0.25 mM ATP. Reactions were carried out at 37 °C for 15 min, and then terminated immediately by addition of the reagent. After incubation of the mixture for 20 min at room temperature, the absorbance at 620 nm was measured using a microplate reader (SpectraMax M2e). The phosphate concentration was determined by calibration with the phosphate standard (BML-KI102).

### Cryo-electron microscopy

To capture different states, the purified Neo1 was mixed with various solutions for 1 h on ice: E1-ATP, 5 mM MgCl_2_ and 2 mM AMPPCP; E2P, 5 mM MgCl_2_, 10 mM NaF, and 2 mM BeSO_4_. After incubation, 2.5-μL aliquots of purified Neo1 at a concentration of about 1 mg/mL were placed on glow-discharged holey carbon grids (Quantifoil Au R2/2, 300 mesh) and were flash-frozen in liquid ethane using a FEI Vitrobot Mark IV. Cryo-EM data was collected automatically with SerialEM in a 300-kV FEI Titan Krios electron microscope with defocus values from −1.0 to −2.0 μm. The microscope was operated with a K3 direct detector at a nominal magnification of 130,000× and a pixel size of 0.413 Å per pixel. The dose rate was 8 electrons per Å^2^ per s, and the total exposure time was 8 s.

### Cryo-EM image processing

Program MotionCorr 2.0 ^1^ was used for motion correction, and CTFFIND 4.1 was used for calculating contrast transfer function parameters ^2^. All the remaining steps were performed using RELION 3 ^3^. The resolution of the map was estimated by the gold-standard Fourier shell correlation at a correlation cutoff value of 0.143.

For the Neo1 structure in the E2P state, we collected 4354 raw movie micrographs. A total of 2,315,630 particles were picked automatically. After 2D classification, a total of 2,068,707 particles were selected and used for 3D classification. Based on the quality of the four 3D classes, 1,279,510 particles were retained for further 3D reconstruction, refinement, and postprocessing, resulting in a 3.25-Å average resolution 3D map.

For the E1-ATP state, we collected 1175 raw movie micrographs. A total of 688,445 particles were picked automatically. After 2D classification, a total of 668,445 particles were selected and used for 3D classification. Based on the quality of the four 3D classes, 264,891 particles were selected for further 3D reconstruction, refinement, and postprocessing, resulting in the 5.64-Å average resolution 3D map.

For the E2P-transition state, we collected 5119 raw movie micrographs. A total of 3,037,926 particles were picked automatically. After 2D classification, a total of 2,661,522 particles were selected and used for 3D classification. Based on the quality of the four 3D classes, 1,673,321 particles were selected for further 3D reconstruction, refinement, and postprocessing, resulting in the 3.08-Å average resolution 3D map.

### Structural modeling, refinement, and validation

We first built the model of Neo1 in the E2P state at 3.2-Å resolution. We generated the initial model based on the structure of Dnf1 (PDB ID 7KYC) by SWISSMODEL (https://swissmodel.expasy.org), and then manually corrected in COOT and Chimera ^4,5^. The complete Neo1 model was refined by real-space refinement in the PHENIX program and subsequently adjusted manually in COOT. Using the model of Neo1 in E2P as reference, models of Neo1 in the E1-ATP and E2P-transition states were built and refined using COOT, Chimera, and PHENIX. Finally, all models were validated using MolProbity in PHENIX ^6,7^. Structural figures were prepared in Chimera and PyMOL (https://pymol.org/2/).

### Yeast strains and plasmid construction

The strains and plasmid used in the study are listed in Supplemental Tables I and II. All yeast culture reagents were purchased from Sigma-Aldrich and BD Scientific, and strains were grown in YPD or minimal selective media. Yeast transformation was performed using the standard LiAC-PEG method ^8^. For plasmid shuffling assays, 10-fold serial dilutions starting from a cell suspension with cell density of OD_600_ = 1 were spotted on synthetic defined media (SD) and SD-5-FOA and incubated at 30 °C or temperatures indicated for at least 2 d before imaging. All images of yeast colonies are representative of three biological replicates (three independently isolated strains of same genotype). DNA constructs and mutations were created by Gibson assembly and quick-change mutagenesis according to the manufacture instructions (using PfuTurbo; Agilent).

### Toxin sensitivity assays

Papuamide A was a kind gift from Raymond Andersen from the University of British Columbia; duramycin was purchased from Sigma Aldrich. For toxin sensitivity assays, mid-log cells were diluted to 0.1 OD_600_ in fresh YPD medium and 100 μL of cells were distributed to each well of 96 well-plate with or without the toxin in 100 μL of YPD. Toxin dilutions were calculated based on final concentrations in a total well volume of 200 μL. Plates were incubated at 30 °C for 20 h. Concentrations of the cells were measured in OD_600_/mL with a Multimode Plate Reader Synergy HT (Biotek). Growth relative to vehicle control (no drug) was used as 100% growth. All values are an average of at least three biological replicates +/− standard deviation.

### Fluorescence microscopy

Strains expressing GFP-Neo1 and Aur1-mKate were grown to mid-log phase in synthetic media. Cells were washed with fresh medium 3 times and resuspended in fresh SD medium. Cells were then mounted on glass slides and observed immediately at room temperature. Images were acquired using a DeltaVision Elite Imaging System (GE Healthcare Life Sciences, Pittsburgh, PA) equipped with a 100X,1.4 NA oil immersion objective lens followed by deconvolution using softWoRx software (GE Healthcare Life Science). Images were analyzed in FIJI (FIJI Is Just ImageJ) using the JaCOP (Just another Colocalization Plugin) add-on. Colocalization was quantified through the computation of the Manders’ coefficient in JaCOP as the fraction of overlap between Neo1 and Aur1.

**Supplementary Information** is available in the online version of the paper.

## ACKNOWLEDGEMENTS

Cryo-EM images were collected in the David Van Andel Advanced Cryo-Electron Microscopy Suite at Van Andel Institute. We thank Gongpu Zhao and Xing Meng for facilitating data collection and David Nadziejka for technical editing of this manuscript. We also thank Raymond Andersen from the University of British Columbia for the papuamide A. This work was supported by the U.S. National Institutes of Health (CA231466 to H.L. and GM107978 to T.R.G.) and the Van Andel Institute (to H.L.).

## CONTRIBUTIONS

L.B., B.K.J., T.R.G., and H.L. conceived and designed the experiments. L.B., B.K.J., Q.Y., and D.D. performed the experiments. L.B., T.R.G., and H.L. analyzed the data. L.B., T.R.G., and H.L. wrote the manuscript with input from all authors.

## DATA AVAILABILITY

The cryo-EM 3D maps and the corresponding atomic models of the Neo1 have been deposited at the EMDB database and the RCSB PDB with the respective accession codes of EMD-xxxx and yyyy (E1P-ATP), EMD-xxxx and yyyy (E2P-transition), EMD-xxxx and yyyy (E2P).

